# Using the aperiodic slope across learning to predict sleep-based memory consolidation

**DOI:** 10.64898/2026.10.02.756363

**Authors:** Annaliese Anesbury, Hayley Bree Caldwell, Kurt Lushington, Maya Al Safadi, Alex Chatburn

## Abstract

Homeostatic mechanisms adjust synaptic strengths to regulate neuronal activity across wake-sleep cycles. Although sleep supports memory consolidation, limited research has explored how sleep-induced homeostatic changes influence memory outcomes in humans. The EEG-derived aperiodic slope (1/*f*) could be a candidate measure of homeostatic memory processes through its reflection of neural excitation-inhibition balance. In this study, 27 participants learned object-word pairs, and completed an immediate recognition test before either a nap or equivalent wakeful period. Resting-state EEG was recorded, both before and after learning and the interval, to derive changes in aperiodic slope values. A delayed recognition test then assessed whether changes in the aperiodic slope (i.e., homeostatic activity) across learning and the interval predicted memory accuracy and N400 amplitude, an index of stimulus expectancy and recognition. Our study found evidence to suggest homeostatic renormalisation, for the nap condition only, indicated by a shift towards inhibition across sleep which was inversely related to the slope change across learning. Although the aperiodic slope change across the nap and wake intervals did not predict delayed memory accuracy, a shift towards inhibition during learning was beneficial for memory accuracy, only in the nap condition. Similarly, a shift towards inhibition across learning, followed by a shift towards excitation in the nap condition, predicted reduced N400 amplitudes. This suggests that homeostatic renormalisation across the nap influenced model updating and stimulus expectancy. This study provides evidence for a link between synaptic homeostatic processes and memory consolidation, emphasising the importance of encoding-related aperiodic activity in this relationship.

## Introduction

Aperiodic activity is a naturally occurring phenomenon within oscillatory systems (Gisiger, 2001). In the brain, aperiodic activity has been previously dismissed as background noise with an assumed lack of functional relevance (He et al., 2010). However, recent research suggests that the aperiodic slope, which reflects the relative low-versus high-frequency power, may provide an index of cortical excitation-inhibition ratio dynamics (i.e., E:I balance), and therefore the homeostatic balance of brain activity (Helfrich et al., 2021). The E:I balance reflects the relative contribution of excitatory and inhibitory synaptic activity within cortical networks and is important for maintaining stable information processing (Vyazovskiy et al., 2009). Following this interpretation, steeper slopes are associated with relatively greater inhibition and flatter slopes with greater excitation (Gao et al., 2017; Vyazovskiy et al., 2008, 2009, 2013).

Across wakefulness, synaptic potentiation increases overall neuronal excitability, which skews the E:I balance to a more excitatory state, incurring cellular energy costs and may reduce the capacity for subsequent learning and memory (Huber et al., 2013; Nere et al., 2013; Vyazovskiy et al., 2009). In contrast, synaptic downscaling during subsequent slow-wave activity (SWA) in non-rapid eye movement (NREM) sleep is proposed to reduce accumulated potentiation and restore homeostatic balance. This process has been shown to promote efficient neural networks, improve signal to noise ratios and support memory consolidation (Huber et al., 2013; Tononi & Cirelli, 2014; Vyazovskiy et al., 2009; Walker & Stickgold, 2005).

It is well known that sleep orchestrates mechanisms that benefit cognitive processes, including learning and memory (Born & Feld, 2012; Rasch & Born, 2013; Walker & Stickgold, 2004; 2006); although, the precise mechanisms underlying this process remain highly debated. The Synaptic Homeostasis Hypothesis (SHY) proposes that the homeostatic regulation of synaptic strength during sleep provides a mechanism through which accumulated synaptic potentiation is reduced, thereby enabling memory consolidation and preserving strongly encoded memories (Tononi & Cirelli, 2003; 2006 2014). In a complementary manner, research suggests that aperiodic activity during sleep may reflect homeostatic E:I balance activity, with the degree of modulation predicting information processing and memory consolidation outcomes (Lender et al., 2022; Helfrich et al., 2021). This implies an important role for aperiodic dynamics in sleep-based memory consolidation. Aperiodic activity may therefore provide a measurable electrophysiological index through which aspects of the theoretical and mechanistic basis of SHY can be investigated. However, whether aperiodic changes over time reflect global homeostatic processes or activity-dependent changes associated with prior learning remains to be tested.

A prospect here is that measuring the aperiodic slope before and after learning, and again across a subsequent nap or period of wakefulness, may help distinguish changes associated with learning from those associated with the retention interval. In this study, we use an object–word associative learning task to examine whether changes in aperiodic slope during learning predict later changes in slope and recognition accuracy. We also test whether slope changes during learning and the subsequent interval relate to N400 amplitude at recognition. Because the N400 is sensitive to the predictability and familiarity of a stimulus (Bornkessel-Schlesewsky et al., 2022; Jano et al., 2024; Hodapp & Rabovsky, 2021), this analysis may indicate whether changes in aperiodic activity are associated with how learned associations are subsequently processed. Together, these measures allow us to examine the relationship between learning-related brain activity, subsequent sleep or wakefulness, and memory performance.

## Methods

### Participants

This study is a reanalysis of a subset of data from Caldwell et al. (2026), using only the wake and nap conditions. Data from 27 individuals was included in the study (25 female, median age =21, range 18-34), with one participant only completing the wake condition before withdrawing, and another participant’s wake condition being excluded due to technical problems (wake *n*=27; nap *n*=27). This sample was based on previous nap studies exploring memory consolidation (Lo et al., 2014; Nishida & Walker, 2007; Payne et al., 2015) Participants completing both conditions did so in different sessions, a minimum of four days apart. Participants were recruited through flyers on the University campus and screened using surveys (see materials) for the following criteria; right-handed, between the ages of 18-35, had no diagnosed psychiatric or sleep conditions, were not taking regular medication that could impact EEG, and had not taken any recreational drugs in the last 6 months had normal or corrected vision and reported no hearing problems, and did not have an extreme chrontype to increase the likelihood of napping (Nicholls et al., 2013; Weymar et al., 2010). Ethical approval for the experiment was acquired from the University of South Australia Human Research Ethics Committee (ID. 205130).

### Materials

#### The Pittsburgh Sleep Quality Index (PSQI)

The PSQI consists of 19 items that are used to calculate self-reported sleep quality over the last month ranging from 0-21 (Buysse et al, 1989). Scores greater than 5 indicate sleep difficulties or poor sleep. Therefore, 5 was used as the cut-off score to determine participant eligibility.

#### Flinders Handedness Survey (Flanders)

The Flanders contains 10 summed items to provide an overall score to screen out participants with left-hand preferences (Nicholls et al., 2013). Participants with scores below 5 were ineligible to participate as this indicated a left-hand preference.

#### Morning-Eveningness Questionnaire (MEQ)

The MEQ (Horne & Östberg, 1976) is a self-report scale querying sleep/wake habits that provides an indication of participant chronotype across 19 questions. Participants who scored in extreme morningness and eveningness categories were not eligible to participate due to circadian factors that may inhibit the likelihood of napping.

#### Karolinska Sleepiness Scale (KSS)

To control for sleepiness as a potential confound The Karolinska Sleepiness Scale was completed by participants to self-report their current level of sleepiness from “1 – extremely alert” to “10 – extremely sleepy, cannot keep awake” (Åkerstedt & Gillberg, 1990).

#### EEG Recording

Participants’ EEG was recorded for resting state, learning, immediate and delayed recognition and sleep phases of the experiment with a 64-channel passive BrainCap with Ag/AgCl electrodes (Brain Products GmbH, Gilching, Germany) positioned according to the modified 10-20 system. Eye movements were captured using electrooculographic electrodes embedded in the cap, positioned 1cm diagonally away from the outer canthus of each eye. Electromyographic activity was measured via three electrodes over the subglottic muscles. The signals were amplified by two BrainAmp amplifiers (DC systems, sampling rate 1000 Hz) and recorded through BrainVision Recorder software with impedances kept below 10kΩ.

#### Stimuli

##### Objects

104 coloured objects were drawn from the Mnemonic Similarity task (MST) with false alarm rates at or below chance level (33.33%; Stark et al., 2019). Within the learning set, each object had a semantically similar counterpart within the set. The object images were formatted at 400 pixels wide, in the centre of the screen.

##### Words

The word stimuli for the pairs were drawn from the Affective Norms for English Words (ANEW), which contains 13,915 English words (Warriner et al., 2013). Words from the ANEW are rated on their frequency in English valence, arousal, and dominance. This study selected 208 monosyllabic nouns (104 per condition) of any frequency rating, but with moderate valence and arousal ratings between 3.5-6.5. The words were presented auditorily with the duration ranging between 500-1000ms each at approximately 70dB, from a recording of a native English speaker with an Australian accent. Objects or words that were highly semantically related within a pair were manually identified and reshuffled.

### Procedure

Participants were instructed to wake up 1 hour earlier than usual on the day of their participation to increase the likelihood of sleep onset and entering NREM in the nap condition. The procedure (see Figure 1) was conducted in the afternoon (12:00-16:30) to capitalise on the natural circadian dip and increase in sleep propensity.

**Figure 1.**
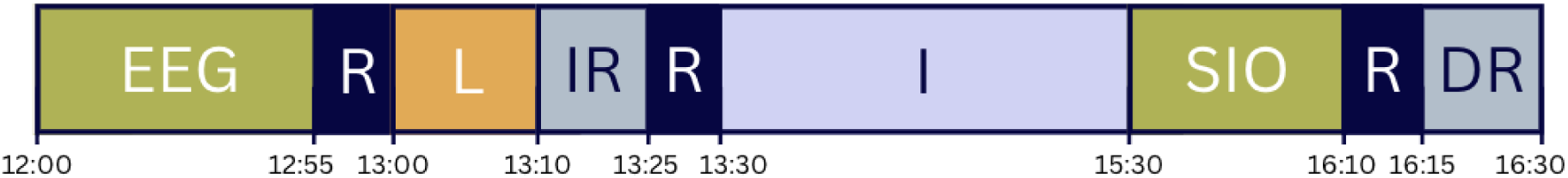
Procedure Timeline. *Note.* EEG = EEG set up, R = Resting State, L = Learning, IR = Immediate Recognition, I = Interval (nap/wake), SIO = Sleep Inertia Offset, DR = Delayed Recognition.

Upon arrival, participants were seated in a room approximately 60cm in front of a computer where they read the study information sheet, signed the consent form, and completed the KSS, FLANDERS and demographic questions. Participants were then fitted with an EEG cap and had their resting state brain activity measured across 2 minutes with eyes closed. Participants first underwent a learning phase, followed by immediate recognition, then another resting state recording just before either a nap or wakeful opportunity. Following a sleep intertia offset period, participants completed a second KSS, resting state and delayed recognition task. The memory paradigm was run in Open Sesame v.3.3.10 (Mathôt et al., 2012; see Figure 2).

**Figure 2.**
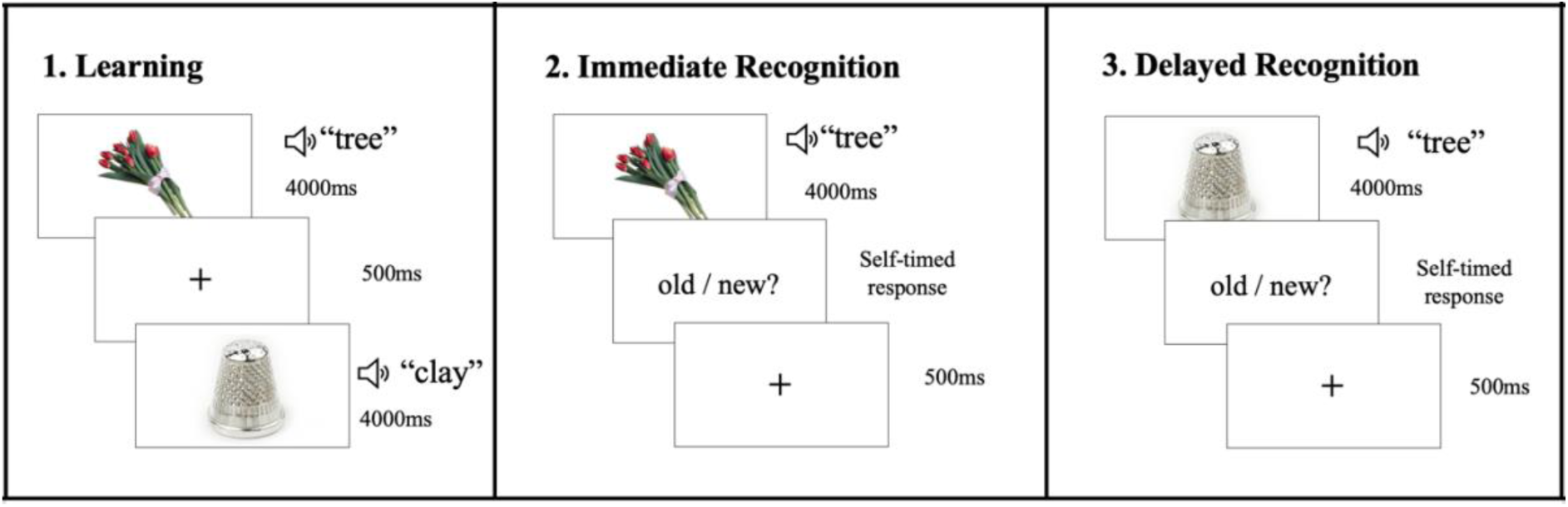
Memory Paradigm. *Note.* 1. The learning phase, participants viewed a object-word pair each trial. 2. In the following task, participants underwent immediate recognition testing where they are required to provide a judgement between original (old) and rearranged (new) pairings. The above is an example of an original (old) pairing as previously seen in the learning phase. 3. The final stage included a delayed recognition task which was the same procedure as immediate recognition with different pairings. The figure depicts an example of a new pairing.

#### Learning

During learning, participants were presented 104 object-word pairs in a randomised order. Objects were presented for 4000ms while the sound of its paired spoken word was played 700ms after the object onset. A 500ms fixation dot (8 pixels wide with a 2-pixel wide centre hole) was presented in between the presentation of each pair. Participants were instructed to learn these objects and words as pairs, with all pairs being presented once. All participants viewed the same object-word pair sets in a randomised order for each condition with a different set of pairs used in each condition.

#### Immediate Recognition

Participants underwent immediate recognition testing using an old/new paradigm. Old pairings were objects and words that were paired in the same way that they were first learnt, whereas new pairings were made up of an object and word from different pairs (i.e., shuffled). object-word pairs were presented in the same way as the learning phase. Following their presentation, the participant was prompted to respond with a keypress if they believed the pair to be old (“Q”) or new (“P”). Participants were encouraged to emphasise accuracy over speed. No feedback was provided on their accuracy.

#### Interval

A second 2-minute resting state with eyes closed was recorded prior to the interval. Participants experienced the experimental (nap) condition, or the control (wake) condition during each of their visits, with the wake condition occurring first to increase environment familiarity and therefore the likelihood of napping.

In the nap condition, participants were provided a 120-minute nap opportunity in a controlled environment; the lights and computer monitor in the room were switched off, white noise was played to suppress distracting sounds, and the participant was asked to turn their phone off. EEG was recorded throughout, with a researcher monitoring for markers of sleep in an adjacent room. Following the nap, participants had a 45-minute period to engage in non-strenuous, self-directed activity, to serve as a sleep inertia offset period.

When participants undertook the wake condition, they were able to engage in tasks of low cognitive demand in the lab for 165 minutes (equivalent to both the 120-minute nap and 45-minute sleep inertia offset period in the nap condition) while maintaining wakefulness. EEG recordings were not taken during the wake period.

5 minutes prior to the conclusion of the break, all participants completed the KSS a second time to measure alertness and undertook a third 2-minute eyes closed resting state period.

#### Delayed Recognition

Participants were presented a similar old/new paradigm as the immediate recognition phase. Participants saw old or new pairings of objects and words, then responded with either old or new judgements. All “new” pairings from the immediate recognition phase were not tested in delayed recognition to avoid interference effects. Following the test, the EEG cap was removed, and the participants received a $200 honorarium after their final session.

### Data Analysis

#### EEG Pre-processing

EEG pre-processing was performed using *MNE-Python* v 1.7.0 (Gramfort et al., 2013). Signals were re-referenced to linked mastoid electrodes (M1 and M2) and down-sampled to 250 Hz. A notch filter (50Hz) removed interference from Australian mains power supply. A zero-phase finite impulse response using a Hamming window of 1-40Hz was applied to remove slow signal drifts and high frequency activity. An Independent Component Analysis (using the *fastica* method) removed data that reflected electrooculographic activity (using the *create_eog_epochs* function in *MNE-Python*). EEG during delayed recognition was segmented into epochs from -200ms pre-word onset and 1000ms post word onset. The *Autoreject* package (Jas et al., 2017) was used to reject bad epochs and repair bad channels using topographic interpolation. Sleep scoring of EEG data was performed in accordance with established procedures from Berry (2012).

#### Aperiodic Slope

Aperiodic slope values were calculated from EEG data obtained from the 2-minutes eyes-closed resting state recordings using the *Irregular-Resampling Auto-Spectral Analysis method* (*IRASA* v 1.0; Wen & Liu, 2016) implemented in the *YASA t*oolbox (Vallat & Walker, 2021). This method separates oscillatory peak components from broadband aperiodic activity (Wen & Liu, 2016). Aperiodic slope values were estimated at three separate timepoints around the experimental phases: pre-learning, pre-interval (i.e., post-learning), and post-interval (see Figure 2). Difference scores of aperiodic slope values were calculated between these timepoints by subtracting the former value form the latter. The learning aperiodic slope change values were the difference score from pre-learning to pre-interval, and interval slope change values were the difference scores from pre-interval to post-interval. These difference scores were calculated per participant, condition and averaged over sagittal (i.e., anterior, central, posterior), and lateral (i.e., left, midline, right) planes to account for topographical variations (Cross; et al., 2024; Jano et al., 2024). Slope change values further from 0 reflect a greater degree of aperiodic slope change, with positive values indicating a shift towards a flatter slope (excitation), and negative values indicating a shift towards a steeper slope (inhibition).

#### N400

N400 ERP activity in the delayed recognition phase was computed in the 1.2 second epochs, across a 350-500ms window following stimulus onset (Hubbard et al., 2019; Jano et al., 2024). Prestimulus activity was calculated in a –200-0 ms window and included in the N400 models as a predictor to control for baseline amplitude differences (Sassenhagen & Alday, 2016). In line with familiarity-based recognition research, fronto-central electrode sites were included in our region of interest (C1, C2, C3, C4, Cz, F1, F2, F3, F4, FC1, FC2, FC3, FC4, Fz) to detect the mean N400 amplitude for hits (correctly identified “old” pairs, indicating familiarity for expected items) per subject and item (Bridger et al., 2012; Stróżak et al., 2016). Fronto-central electrodes, as above, were used when plotting the grand average, using the *ggplot2* package in *R* v 4.1.3 (Wickham, 2016).

#### Recognition Accuracy

We captured recognition accuracy with *d-*prime *(d’)* calculated in *R* v 4.1.3 (R Core Team, 2022) using the *Psycho* package (Makowski, 2018). The *d’* values were calculated as the difference between the z-transformed probabilities of hit rate (HR) and false alarm rate (FA; i.e., *d’* = z[HR] – z[FA]), and was calculated. Recognition accuracy *d’* was calculated per participant for their judgements of pairings as old or new in the immediate recognition and delayed recognition phase.

#### Statistical Analysis

To test various predictions, five linear mixed-effects models (LMM) fit by Restricted Maximum Likelihood were conducted in *R* v 4.1.3 (R Core Team, 2022) using the *lmerTest* package (Kuznetsova et al., 2017). Outliers were identified and removed for each dependent variable. Observations falling more than 1.5 interquartile ranges below the first quartile or above the third quartile were classified as outliers. Sum contrasts were applied to all categorical variables condition [nap, wake], sagittal [anterior, central, posterior] and lateral planes [left, midline, right] (with the *contr.Sum* function from the *car* package), to assess the deviation of each level from the grand mean, represented by coefficients (Brehm & Alday, 2022; Fox & Weisberg, 2019). Statistical significance for fixed effects was calculated via type II Wald chi-square tests from the *Anova* function in the *car* package (Fox & Weisberg, 2019). Significant main and interaction effects from the linear mixed models were derived using *effects* (Fox & Weisberg, 2019) and plotted with *ggplot2* (Wickham, 2016). The error bars were set to 83% confidence intervals, where non-overlapping error estimates reflect 5% significance thresholds (Austin & Hux, 2002; MacGregor-Fors & Payton, 2013).

The first model tested whether learning slope change values predicts interval slope changes across conditions. The outcome variable was interval slope changes, predicted by the fixed effects of learning slope change, condition (nap or wake), sagittal regions, and lateral regions to account for topographical differences. Scores of the first KSS were to control for sleepiness as a suggested explanation for aperiodic slope changes (Chatburn et al., 2024). Finally, subject ID was used as a random effect:

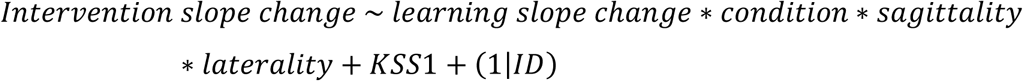

The second model tested whether the interval slope change and condition (i.e., nap or wake) predicted delayed memory recognition. The outcome variable was delayed recognition *d’* scores, predicted by the fixed effect of condition. Immediate recognition *d’* scores and scores of the second KSS were covariates, with subject ID as a random intercept:

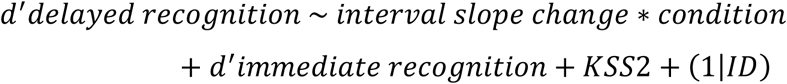

The third model tested if aperiodic learning slope change and condition predicts delayed memory recognition. The outcome variable was delayed recognition *d’* scores predicted by fixed effects of aperiodic slope change across learning and between conditions. Scores of the second KSS and immediate recognition *d’*, was included as covariates with subject ID as the random intercept:

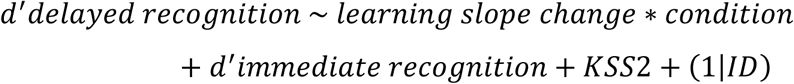

The fourth model tested the potential interaction between learning slope changes and interval slope changes, along with condition to predict delayed memory recognition. The outcome variable was delayed recognition *d’* scores predicted by fixed effects of learning aperiodic slope change, interval aperiodic slope change and condition. Scores of the second KSS and immediate recognition *d’* were included as covariates with subject ID as the random intercept:

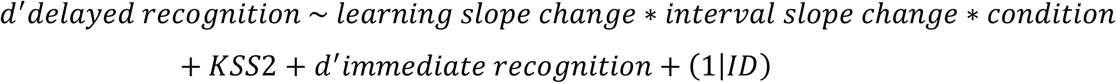

The final model aimed to investigate the functional implications of aperiodic slope change and recognition-related neural activity (N400). The outcome variable was delayed recognition N400 amplitude for hits (i.e., old pairings corerectly identified as old), predicted by fixed effects of learning and interval slope change and condition. Mean N400 pre-stimulus activity was included as a covariate and subject ID as a random intercept. The model is outlined as:

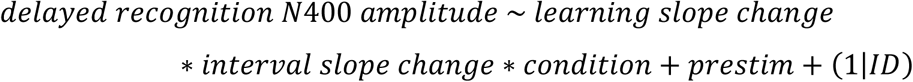

## Results

Table 1 summarises mean *d*’ scores for each experimental phase, and total overall scores. All participants scored above chance.

**Table 1.** Recognition Accuracy *d’* Scores Across Testing Phases and Conditions.

| Interval Condition | Testing Phase |  |
| --- | --- | --- |
| | Mean Immediate Recognition<br>$d'$ (SD) | Mean Delayed<br>Recognition $d'$ (SD) |
| Nap | 1.01 (0.77) | 1.42 (0.91) |
| Wake | 0.75 (0.50) | 0.94 (0.59) |
| Total | 0.88 (0.65) | 1.18 (0.79) |

*Note.* Mean values are reported across participants, between conditions. SD=standard deviation, reported in brackets.

Table 2 shows the sleep parameters from the EEG recordings during the nap interval. All participants slept during the designated nap period, and experienced NREM sleep, with only five subjects not experiencing slow-wave sleep. Four subjects entered rapid eye movement sleep during the nap.

**Table 2.** Sleep Parameter Statistics.

| Sleep Parameters | Mean (SD) | Range |
| --- | --- | --- |
| TST (mins) | 82.44 (26.31) | 25.00 – 116.00 |
| SOL (mins) | 11.90 (11.11) | 1.50 – 53.00 |
| N1 (%) | 16.70 (9.54) | 5.00 – 49.50 |
| N2 (%) | 43.90 (14.00) | 19.50 – 71.00 |
| SWS (%) | 19.00 (18.31) | 0.00 – 56.00 |
| REM (%) | 2.87 (6.30) | 0.00 – 20.50 |

*Note:* TST = total sleep time, SOL = sleep onset latency, N1 = stage 1 NREM sleep, N2 = stage 2 NREM sleep, SWS = slow wave sleep, REM = rapid eye movement sleep.

### Shifts in learning slope change predict interval slope changes for the nap condition

To test if changes in the aperiodic slope across learning were related to subsequent slope changes across the nap, the first LMM was conducted to predict their relationship, by condition. This model revealed a significant interaction of learning slope change and condition to predict interval slope change, χ^2^(1) = 15.20, *p* < .001. A flatter (more positive) slope during learning was associated with a shift towards a steeper (more negative) slope across the interval, which was significantly more pronounced in the nap condition, relative to the grand mean (*B* = –0.20, *SE* = 0.05, *p* < .001) (see supplementary materials for full model output).

**Figure 3.**
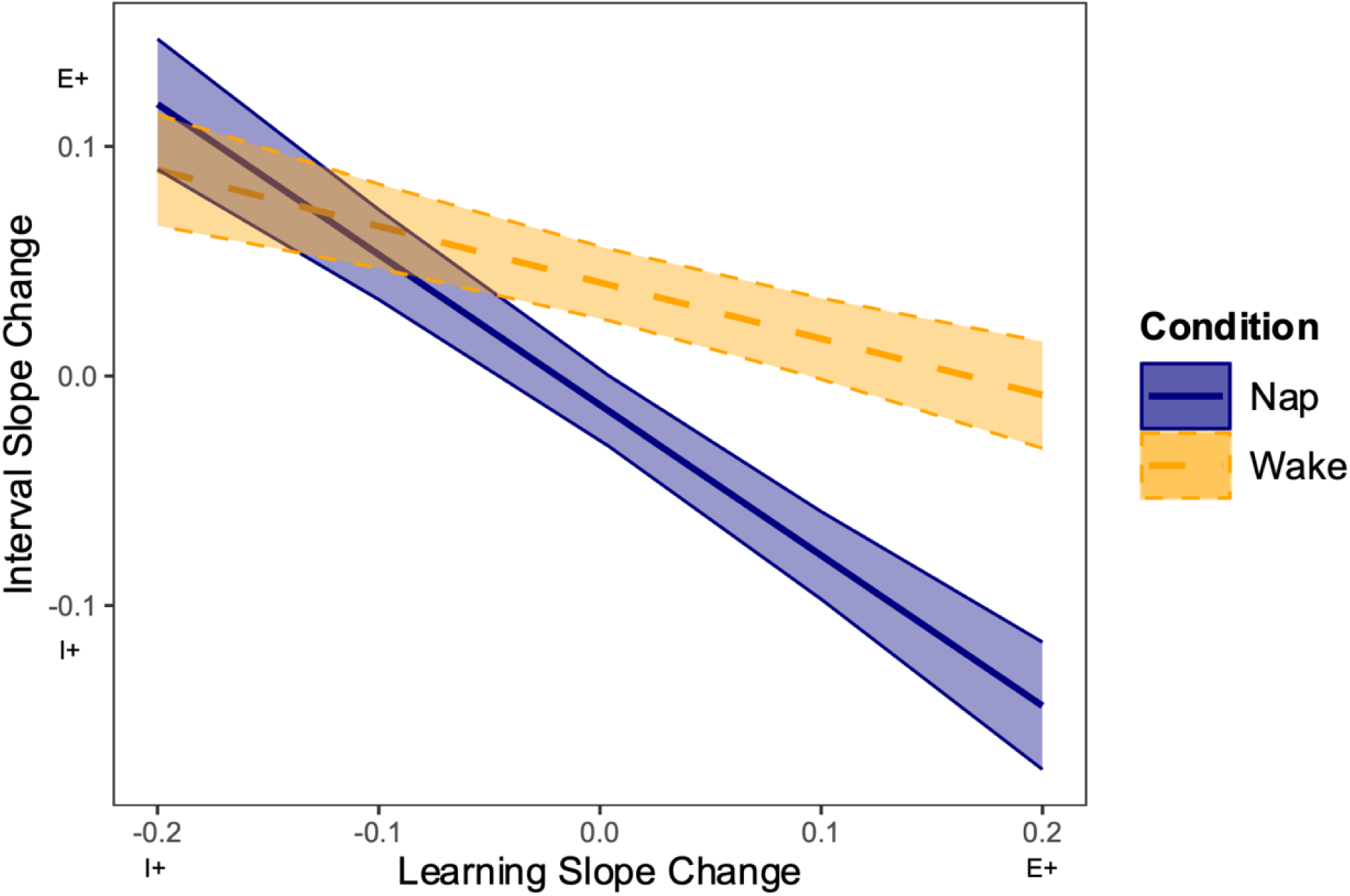
Aperiodic Slope Change Across Interval and Learning. *Note.* Interval slope change (y-axis) by learning slope change (x-axis) and condition (nap = purple solid line, wake = orange dashed line). Shaded regions reflect 83% confidence intervals. Slope change values further from 0 reflect a greater slope change, with positive values indicating a shift towards a flatter slope (E+; excitation), and negative values indicating a shift towards a steeper slope (I+; inhibition).

### Learning aperiodic slope changes predict delayed recognition accuracy for the nap condition

A second LMM was conducted to predict delayed recognition accuracy from interval slope change values and condition. This model showed a main effect of condition on delayed recognition accuracy χ^2^(1) = 36.19, *p* < .001. However, the model revealed no significant interaction effects for interval slope change on delayed recognition accuracy between conditions, χ^2^(1) = 3.04, *p* =.08.

The third LMM examined whether learning slope changes and condition predicted delayed recognition accuracy. This model revealed a significant interaction between learning slope change and condition to predict delayed recognition accuracy, χ^2^(1) = 5.12, *p* = .02. Figure 4 depicts that a shift towards a steeper (more negative) slope in learning, resulted in better memory accuracy at testing, but only for the nap condition, *B* = –0.31, *SE* = 0.14, *p* = .02.

**Figure 4.**
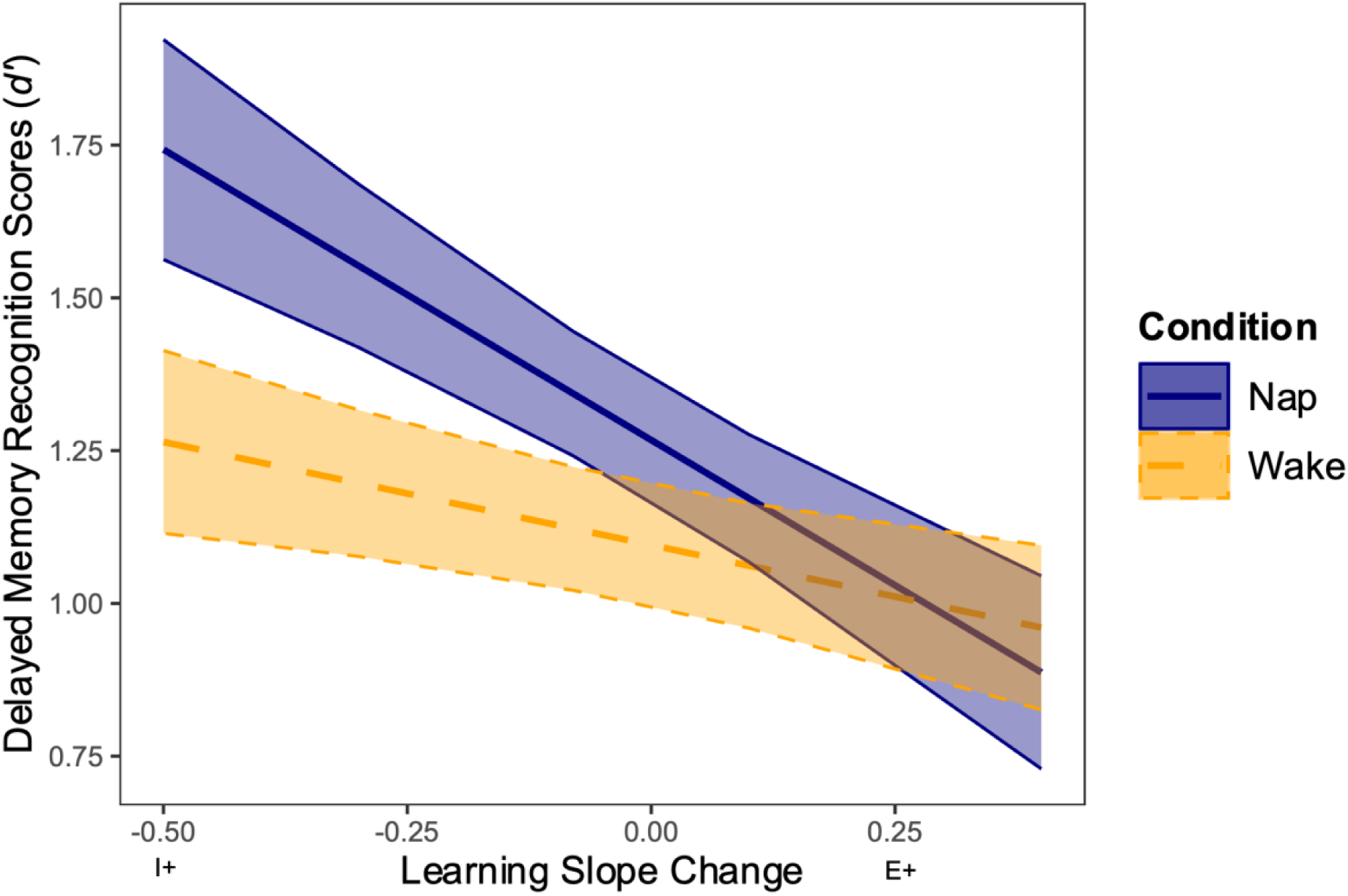
Aperiodic slope change over learning on memory at delayed recognition between conditions. *Note. d*-prime memory recognition accuracy (y-axis) by learning slope change (x-axis) and condition (nap = purple solid line, wake = orange dashed line). Shaded regions reflect 83% confidence intervals. Slope change values further from 0 reflect a greater slope change, with positive values indicating a shift towards a flatter slope (E+; excitation), and negative values indicating a shift towards a steeper slope (I+; inhibition).

Lastly, the fourth LMM explored the interactive process that learning slope changes, interval slope changes and condition may have on delayed memory recognition.This model did not reveal a significant interaction effect between learning slope change, interval slope change and condition on delayed memory recognition, χ^2^(1) = 0.01, *p* = .92, but did support a main effect of learning slope change on delayed memory recognition (χ*^2^*(1) = 11.12, *p* < .001; see Figure 5).This main effect demonstrates that learning slope change was negatively related to delayed recognition scores (similarly to Figure 4) regardless of condition.

### Functional implications of aperiodic slope changes and memory accuracy

To explore the functional implications of changes in the aperiodic slope on memory, a fifth LMM was conducted using N400 amplitudes. First, a grand average ERP was plotted across fronto-central electrodes between nap and wake conditions, which demonstrates a negative deflection around 400ms (see Figure 6a) This model revealed a significant interaction between learning slope change, interval slope change and condition, which predicted delayed recognition N400 amplitude, χ^2^(1)=3.97, *p* = .046.

**Figure 6.**
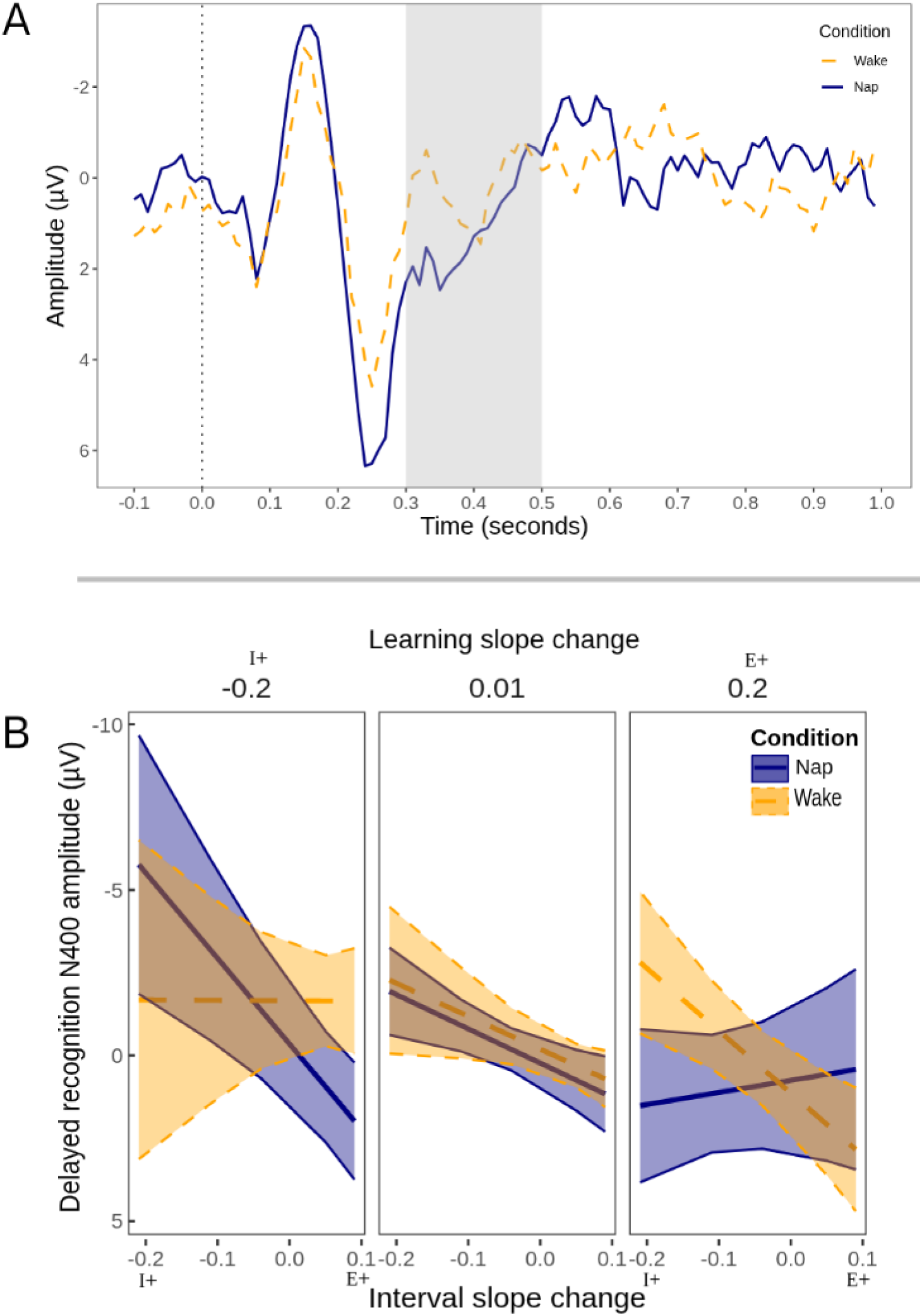
Aperiodic slope change across learning and interval phases predicting N400 amplitude by condition. *Note:* A. Grand average ERP component at fronto-central electrodes. The grey shaded area represents the time window (300-500ms), of the N400 amplitude plotted on the y-axis by condition (wake = orange dashed line, nap = purple solid line), with negativity plotted upwards. B. The modelled results of the interval slope change (y-axes) by learning slope change (facets) and condition (wake = orange dashed line, nap = purple solid line). Shaded regions reflect 83% confidence intervals, where the non-overlapping line indicates significance. Slope change values further from 0 reflect a greater slope change, with positive values indicating a shift towards a flatter slope (E+; excitation), and negative values indicating a shift towards a steeper slope (I+; inhibition). Negative values indicate greater N400 amplitudes and positive values indicate smaller N400 amplitudes. Facets were grouped via the output of the modelled effects for representative values across the observed range of learning and interval slope changes (from –0.20 to 0.20).

Figure 6b depicts that a shift towards a steeper slope (i.e., inhibition; more negative) in learning, coupled with a shift towards a flatter slope (i.e., excitation; more positive) across the interval, related to reduced (less negative) N400 amplitudes, but only for the nap condition *B* = –60.21, *SE* = 30.20, *p* = .047.

## Discussion

This study investigated how changes in the aperiodic slope relate to learning and memory across sleep versus wakefulness, through a homeostatic lens. Our results suggested a relationship between the learning slope change and the interval slope change, whereby flatter, excitatory shifts across learning are subsequently renormalised through steeper, inhibitory shifts across sleep, but not wake. This finding supports theories of homeostatic downscaling across sleep (Huber et al., 2013; Lendner et al., 2022). The interval slope change did not impact memory outcomes, either in consideration of learning slope changes or in isolation. However, in contrast to the pattern above, a shift towards a steeper slope in learning was found to better relate to memory accuracy, only if learning was followed by a nap. Further, ERP results showed that a shift towards a steeper slope across learning, coupled with a shift towards a flatter slope following sleep, was associated with reduced N400 amplitudes. Therefore, a shift towards inhibition across learning followed by excitation across subsequent sleep may provide the best conditions for effective model updating, reflected by reduced expectancy violations for correctly presented pairs. These findings support the idea that aperiodic changes from learning to across the interval are necessary for neurophysiological processes but are not entirely necessary for behavioural accuracy, whereas aperiodic changes across encoding, coupled with a nap, are most important to demonstrate greater memory performance outcomes.

Our findings of a net shift towards excitation during learning accompanied by a corresponding shift towards inhibition across sleep support recent research into changes in aperiodic networks across sleep stages and synaptic homeostasis (Lendner, et al., 2022; Niethard et al., 2017; Olcese et al., 2010). NREM sleep typically dominates an afternoon nap period, and SWA within NREM reflects homeostatic pressure to support synaptic renormalisation (Tononi, 2009; Walker & Stickgold, 2006). These NREM attributes offer a plausible mechanism of homeostasis that could underlie the aperiodic changes observed here. Consistent with this interpretation, Helfrich et al. (2021) proposed a role for aperiodic activity, interleaved with SWA, in imprinting new mnemonic patterns onto existing schemas. Although their account primarily focuses on the interplay between aperiodic activity within sleep and SWA for memory consolidation, it motivates the question of whether aperiodic activity occurring in tandem with SWA also contributes to broader changes in the overall aperiodic slope, as suggested by Lendner et al. (2022). At present, it is unclear whether the mechanisms responsible for sleep-related aperiodic change reflect local processes supporting memory consolidation, broader homeostatic changes in E:I balance, or both.

In addition to the canonical pattern of a renormalisation of aperiodic activity across a nap, our study also observed an inverse pattern; a shift towards inhibition across learning resulted in a shift towards excitation across the interval, irrespective of whether the interval involved a nap or wakefulness. One interpretation of this inverse relationship is that it reflects a homeostatic response to learning-specific activity, though the underlying basis likely differs by state. During a nap, this reversal may function to restore E:I balance by upregulating excitatory networks or downscaling inhibitory synaptic strengths (Tononi & Cirelli, 2012), whereas during wake, it likely reflects the natural accumulation of excitatory drive with increasing time awake (Borbely et al., 2016). Previous work has predominately focused on the renormalisation of net excitatory increases accumulated throughout the day (Vyazovskiy et al., 2008). Our findings align with these results and additionally extend them by demonstrating that sleep may also renormalise shifts in inhibitory activity arising during wake (Tononi & Cirelli, 2014). However, as the aperiodic slope reflects the ratio of the E:I balance, an apparent shift towards inhibition may reflect either increased inhibitory activity or a reduction in excitatory activity, and vice versa. These changes are therefore best understood as reflecting dynamic modulation of overall E:I balance, rather than evidence for excitatory or inhibitory activity in isolation.

Contrary to our predictions, changes in aperiodic slope across the interval did not predict subsequent memory performance. However, inhibitory shifts in the aperiodic slope across learning, predicted increased subsequent memory performance for the nap condition relative to the wake condition. This suggests that encoding-related E:I dynamics are important for providing the optimal environment for sleep-based memory consolidation. One interpretation is that a shift towards inhibitory activity during encoding may reflect a stabilisation of network activity with increased signal-to-noise ratios (Niethard et al., 2017). Inhibitory processes have been implicated in regulating information processing and neuronal plasticity, protecting newly encoded memories from interference and stabilising neural networks to promote consolidation (Barron, 2021; Litwin-Kumar & Doiron, 2014; Niethard et al., 2017; Ognjanovski et al., 2017). Our findings extend this by suggesting that broader network-level encoding-related changes may facilitate newly formed associative representations amenable to sleep-based memory consolidation. Whilst optimal encoding states within frequency bands have been established (e.g., theta power; Gibson et al., 2022), these findings are the first to suggest aperiodic states may also contribute to an facilitatory encoding environment for sleep-based memory consolidation. Further, these findings provide more evidence to suggest that sleep-based memory consolidation does not occur in isolation, and factors across the frequency spectrum during learning should be taken into consideration when exploring the mechanisms and outcomes of sleep-based memory consolidation.

Our findings supported the prediction that reduced N400 amplitudes reflect memories being incorporated into internal models, under certain aperiodic changes across learning and sleep. The N400 typically indexes expectancy violation and internal model updating (Hodapp & Rabovsky, 2021; Kuperberg et al., 2020; Rabovsky et al., 2018). Therefore, our results may indicate that expectancy violation was significantly reduced for correctly identified object-word pairings at delayed recognition as they were perceived as congruent with mental models. Specifically, we observed the largest reduction in expectancy violation when learning showed a shift towards inhibition, followed by a corresponding excitatory shift across sleep. Consistent with our behavioural accuracy results, a shift towards inhibitory activity during learning likely reflects an improved signal-to-noise ratio environment. This reduction in neural noise is conducive to the formation moderately efficient models, which is more preferentially acted upon by sleep-based memory consolidation (Niethard et al., 2017; Stickgold, 2009). Together, this improved model updating reflects the result of complementary sleep-based aperiodic changes and optimal encoding conditions (Tononi & Cirelli, 2014). Our results therefore support that the homeostatic renormalisation of E:I balance across post-learning sleep may support enhanced model updating under optimal encoding conditions, which could implicate homeostasis as a key process for the refinement of models.

While these results highlight an important role for the aperiodic slope in reflecting state-like fluctuations across time and tasks, prior work suggests that the aperiodic slope also reflects trait-like individual differences in information processing, recognition tasks, and language processing (Bornkessel-Schlesewsky et al., 2022; Cross et al., 2022; Dave et al., 2018; Donoghue et al., 2020; Immink et al., 2021; Ouyang et al., 2020; Voytek & Knight, 2015). One interpretation of the present results in light of this could be that there are trait-like variations in aperiodic activity, which interact with state-dependent fluctuations, perhaps in a rank order. Disentangling the contributions of trait- and state-like aperiodic activity would require repeated sampling across multiple sessions, with resting state recordings independent of those used to derive state-like changes. Future longitudinal work should therefore examine whether state-like fluctuations in the aperiodic slope are moderated by stable, trait-like individual differences in baseline aperiodic slope.

Overall, our findings suggest that aperiodic activity during encoding is an important facet of behavioural and neural memory outcomes, extending its role beyond sleep-related changes in global network activity. A shift towards inhibition during learning was associated with better behavioural memory accuracy following sleep. The subsequent homeostatic reversal across sleep, though not independently predictive of behavioural accuracy, was associated with electrophysiological evidence of model updating, as indexed by the N400. These findings suggest a link between synaptic homeostasis and memory outcomes and highlight the role of aperiodic activity in indexing this process.

## Supplementary Materials

**Model 1 output:**
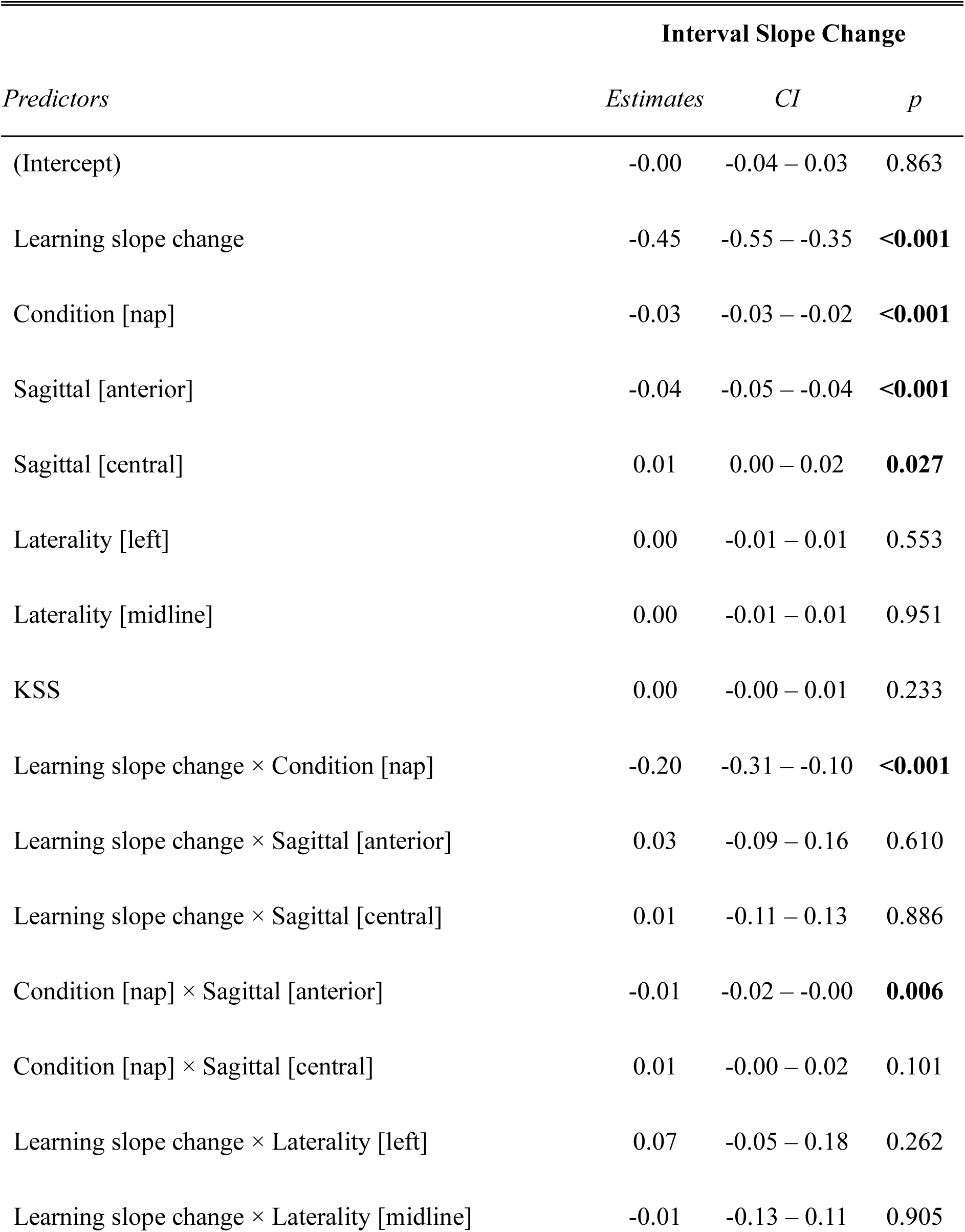
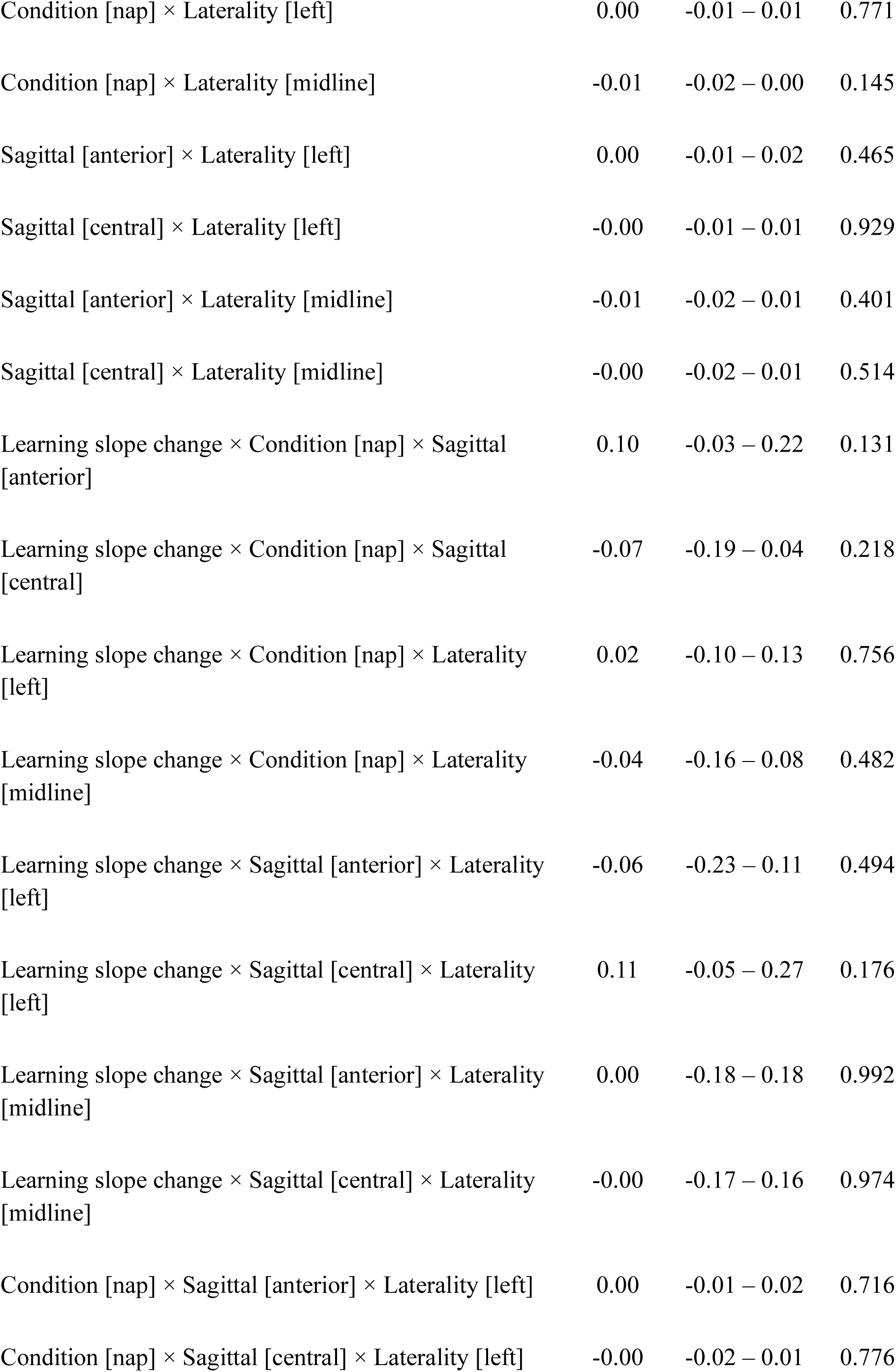
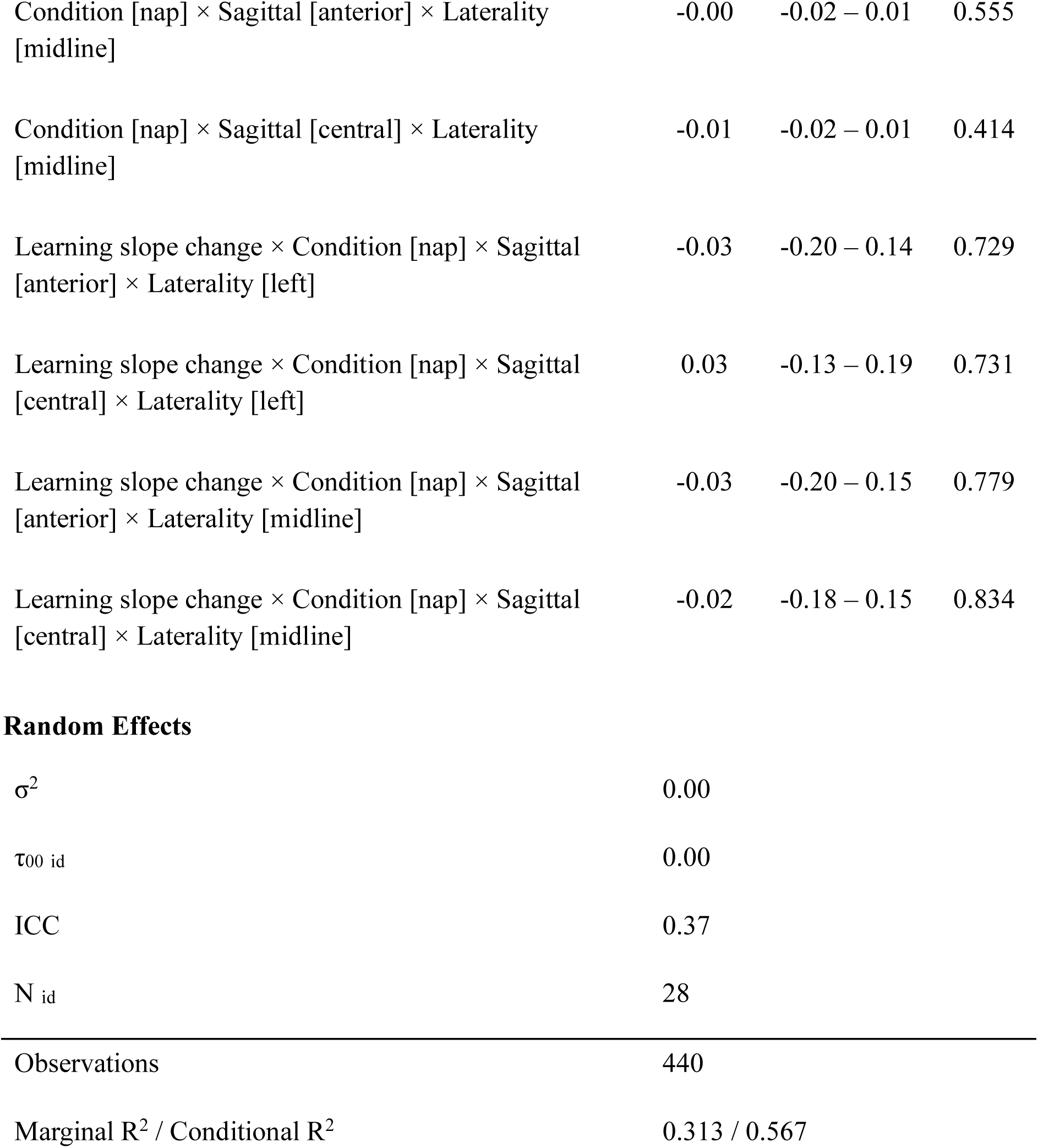
Learning slope change predicting interval slope change by condition.

| *Predictors* | Interval Slope Change |  |  |
| --- | --- | --- | --- |
|  | *Estimates* | *CI* | *p* |
| (Intercept) | -0.00 | -0.04 – 0.03 | 0.863 |
| Learning slope change | -0.45 | -0.55 – -0.35 | **<0.001** |
| Condition [nap] | -0.03 | -0.03 – -0.02 | **<0.001** |
| Sagittal [anterior] | -0.04 | -0.05 – -0.04 | **<0.001** |
| Sagittal [central] | 0.01 | 0.00 – 0.02 | **0.027** |
| Laterality [left] | 0.00 | -0.01 – 0.01 | 0.553 |
| Laterality [midline] | 0.00 | -0.01 – 0.01 | 0.951 |
| KSS | 0.00 | -0.00 – 0.01 | 0.233 |
| Learning slope change × Condition [nap] | -0.20 | -0.31 – -0.10 | **<0.001** |
| Learning slope change × Sagittal [anterior] | 0.03 | -0.09 – 0.16 | 0.610 |
| Learning slope change × Sagittal [central] | 0.01 | -0.11 – 0.13 | 0.886 |
| Condition [nap] × Sagittal [anterior] | -0.01 | -0.02 – -0.00 | **0.006** |
| Condition [nap] × Sagittal [central] | 0.01 | -0.00 – 0.02 | 0.101 |
| Learning slope change × Laterality [left] | 0.07 | -0.05 – 0.18 | 0.262 |
| Learning slope change × Laterality [midline] | -0.01 | -0.13 – 0.11 | 0.905 |
| Condition [nap] × Laterality [left] | 0.00 | -0.01 – 0.01 | 0.771 |
| Condition [nap] × Laterality [midline] | -0.01 | -0.02 – 0.00 | 0.145 |
| Sagittal [anterior] × Laterality [left] | 0.00 | -0.01 – 0.02 | 0.465 |
| Sagittal [central] × Laterality [left] | -0.00 | -0.01 – 0.01 | 0.929 |
| Sagittal [anterior] × Laterality [midline] | -0.01 | -0.02 – 0.01 | 0.401 |
| Sagittal [central] × Laterality [midline] | -0.00 | -0.02 – 0.01 | 0.514 |
| Learning slope change × Condition [nap] × Sagittal [anterior] | 0.10 | -0.03 – 0.22 | 0.131 |
| Learning slope change × Condition [nap] × Sagittal [central] | -0.07 | -0.19 – 0.04 | 0.218 |
| Learning slope change × Condition [nap] × Laterality [left] | 0.02 | -0.10 – 0.13 | 0.756 |
| Learning slope change × Condition [nap] × Laterality [midline] | -0.04 | -0.16 – 0.08 | 0.482 |
| Learning slope change × Sagittal [anterior] × Laterality [left] | -0.06 | -0.23 – 0.11 | 0.494 |
| Learning slope change × Sagittal [central] × Laterality [left] | 0.11 | -0.05 – 0.27 | 0.176 |
| Learning slope change × Sagittal [anterior] × Laterality [midline] | 0.00 | -0.18 – 0.18 | 0.992 |
| Learning slope change × Sagittal [central] × Laterality [midline] | -0.00 | -0.17 – 0.16 | 0.974 |
| Condition [nap] × Sagittal [anterior] × Laterality [left] | 0.00 | -0.01 – 0.02 | 0.716 |
| Condition [nap] × Sagittal [central] × Laterality [left] | -0.00 | -0.02 – 0.01 | 0.776 |
| Condition [nap] × Sagittal [anterior] × Laterality [midline] | -0.00 | -0.02 – 0.01 | 0.555 |
| Condition [nap] × Sagittal [central] × Laterality [midline] | -0.01 | -0.02 – 0.01 | 0.414 |
| Learning slope change × Condition [nap] × Sagittal [anterior] × Laterality [left] | -0.03 | -0.20 – 0.14 | 0.729 |
| Learning slope change × Condition [nap] × Sagittal [central] × Laterality [left] | 0.03 | -0.13 – 0.19 | 0.731 |
| Learning slope change × Condition [nap] × Sagittal [anterior] × Laterality [midline] | -0.03 | -0.20 – 0.15 | 0.779 |
| Learning slope change × Condition [nap] × Sagittal [central] × Laterality [midline] | -0.02 | -0.18 – 0.15 | 0.834 |

Random Effects
|  |  |
| --- | --- |
| $\sigma^2$ | 0.00 |
| $\tau_{00 \text{ id}}$ | 0.00 |
| ICC | 0.37 |
| $N_{\text{id}}$ | 28 |
| Observations | 440 |
| Marginal $R^2$ / Conditional $R^2$ | 0.313 / 0.567 |

**Model 2 output:** interval slope change predicting delayed recognition *d’* by condition.

| Delayed recognition $d'$ | | | |
| --- | --- | --- | --- |
| *Predictors* | *Estimates* | *CI* | *p* |
| (Intercept) | 0.82 | 0.63 – 1.00 | <0.001 |
| Interval slope change | 0.05 | -0.19 – 0.30 | 0.675 |
| Condition [nap] | 0.09 | 0.06 – 0.12 | <0.001 |
| Immediate recognition $d'$ | 0.77 | 0.70 – 0.84 | <0.001 |
| KSS | -0.07 | -0.09 – -0.05 | <0.001 |
| Interval slope change × Condition [nap] | 0.19 | -0.02 – 0.41 | 0.082 |
| **Random Effects** |  |  |  |
| $\sigma^2$ | 0.06 | | |
| $\tau_{00 \text{ id}}$ | 0.13 | | |
| ICC | 0.67 |  |  |
| N<sub>id</sub> | 28 |  |  |
| Observations | 477 |  |  |
| Marginal $R^2$ / Conditional $R^2$ | 0.593 / 0.866 | | |

**Model 3 output:** learning slope change predicting delayed recognition *d’* by condition.

| Delayed recognition $d'$ | | | |
| --- | --- | --- | --- |
| *Predictors* | *Estimates* | *CI* | *p* |
| (Intercept) | 0.84 | 0.66 – 1.03 | <0.001 |
| Learning slope change | -0.64 | -0.92 – -0.37 | <0.001 |
| Condition [nap] | 0.09 | 0.06 – 0.11 | <0.001 |
| KSS | -0.08 | -0.10 – -0.05 | <0.001 |
| Immediate recognition $d'$ | 0.75 | 0.68 – 0.83 | <0.001 |
| Learning slope change × Condition [nap] | -0.31 | -0.57 – -0.04 | 0.024 |
| **Random Effects** |  |  |  |
| $\sigma^2$ | 0.06 | | |
| $\tau_{00 \text{ id}}$ | 0.14 | | |
| ICC | 0.69 |  |  |
| N<sub>id</sub> | 28 |  |  |
| Observations | 477 |  |  |
| Marginal $R^2$ / Conditional $R^2$ | 0.592 / 0.873 | | |

**Model 4 output:**
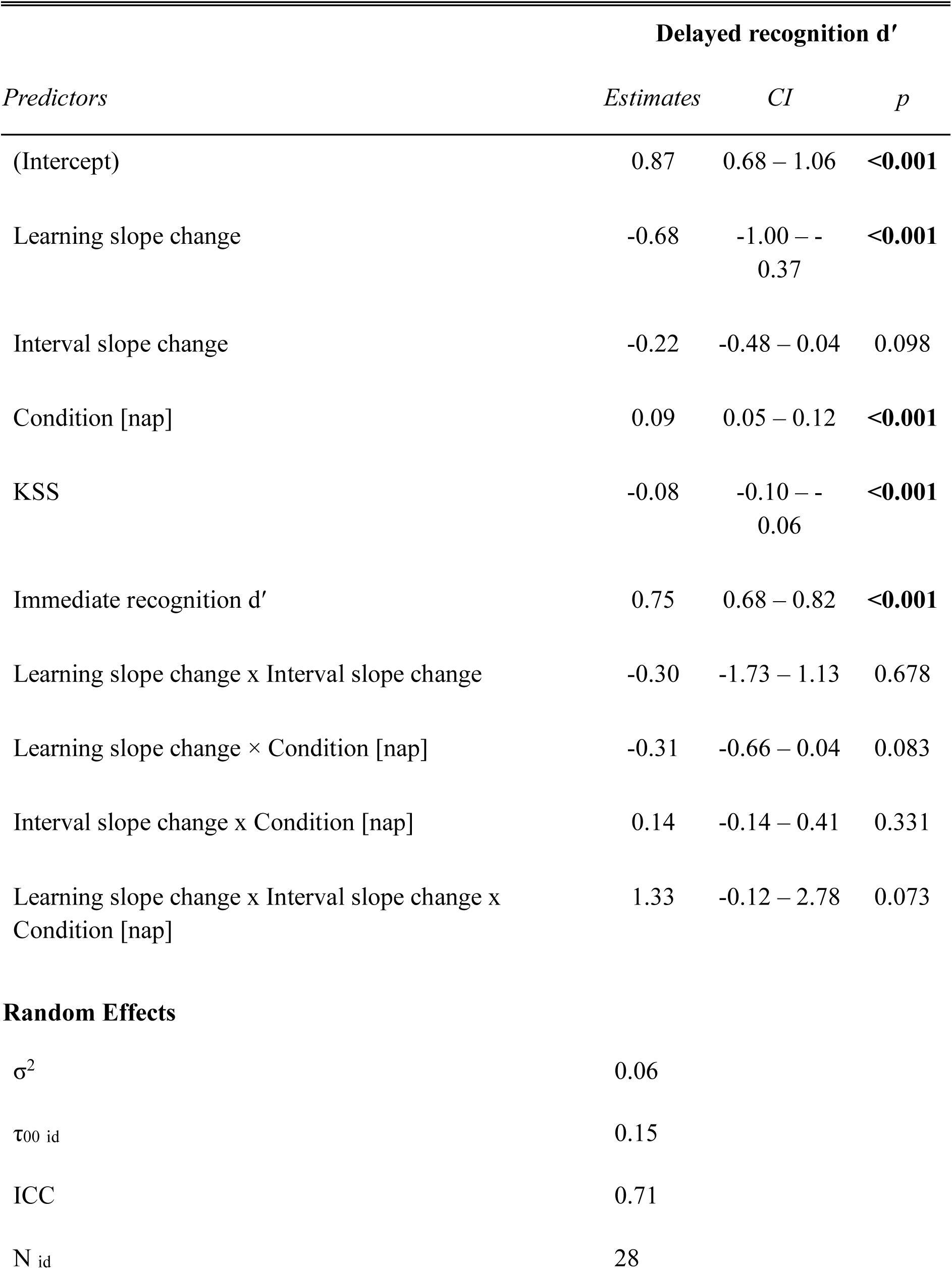
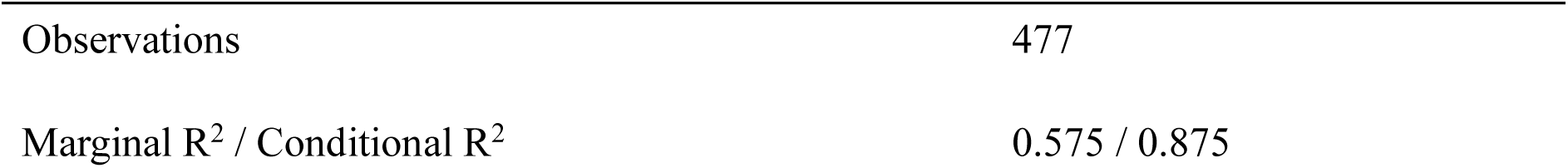
learning slope change and interval slope change predicting delayed recognition *d’* by condition.

| *Predictors* | **Delayed recognition $d'$** |  |  |
| --- | --- | --- | --- |
|  | *Estimates* | *CI* | *p* |
| (Intercept) | 0.87 | 0.68 – 1.06 | **<0.001** |
| Learning slope change | -0.68 | -1.00 – -<br>0.37 | **<0.001** |
| Interval slope change | -0.22 | -0.48 – 0.04 | 0.098 |
| Condition [nap] | 0.09 | 0.05 – 0.12 | **<0.001** |
| KSS | -0.08 | -0.10 – -<br>0.06 | **<0.001** |
| Immediate recognition $d'$ | 0.75 | 0.68 – 0.82 | **<0.001** |
| Learning slope change x Interval slope change | -0.30 | -1.73 – 1.13 | 0.678 |
| Learning slope change × Condition [nap] | -0.31 | -0.66 – 0.04 | 0.083 |
| Interval slope change x Condition [nap] | 0.14 | -0.14 – 0.41 | 0.331 |
| Learning slope change x Interval slope change x<br>Condition [nap] | 1.33 | -0.12 – 2.78 | 0.073 |
| **Random Effects** |  |  |  |
| $\sigma^2$ | 0.06 | | |
| $\tau_{00 \text{ id}}$ | 0.15 | | |
| ICC | 0.71 |  |  |
| $N_{\text{id}}$ | 28 | | |

|  |  |
| --- | --- |
| Observations | 477 |
| Marginal $R^2$ / Conditional $R^2$ | 0.575 / 0.875 |

**Model 5 output:**
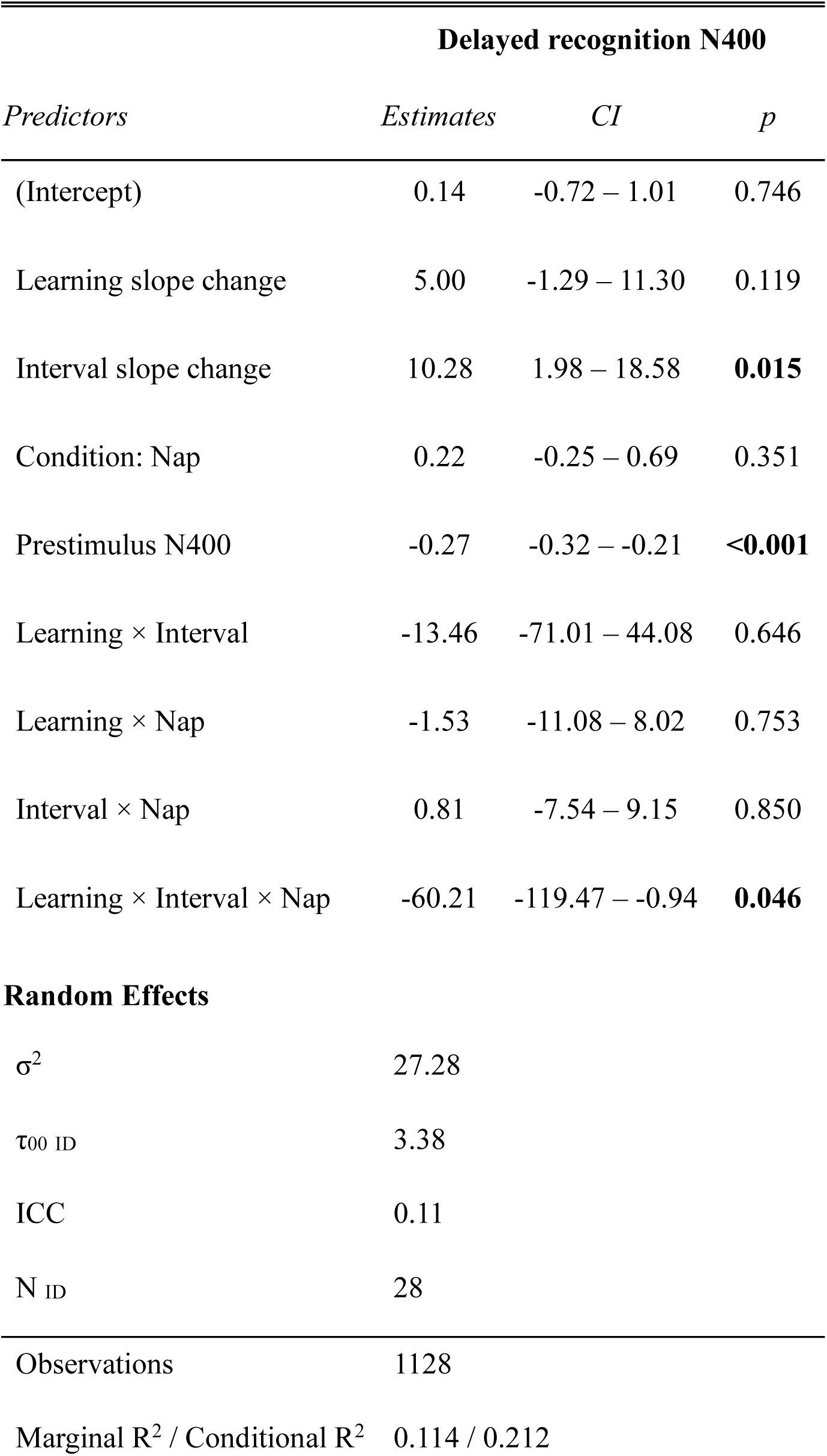
learning slope change and interval slope change predicting delayed recognition N400 amplitude by condition.

## Notes

### Competing Interest Statement

The authors have declared no competing interest.

## References

Akerstedt, T., & Gillberg, M. (1990). Subjective and objective sleepiness in the active individual. The International Journal of Neuroscience, 52(1–2), 29–37. 10.3109/00207459008994241

Austin, P. C., & Hux, J. E. (2002). A brief note on overlapping confidence intervals. Journal of Vascular Surgery, 36(1), 194–195. 10.1067/mva.2002.125015

Barron, H. C. (2021). Neural inhibition for continual learning and memory. Current Opinion in Neurobiology, 67, 85–94. 10.1016/j.conb.2020.09.007

Berry, R. B. (2012). Chapter 3—Sleep staging in adults. In R. B. Berry (Ed.), Fundamentals of sleep medicine (pp. 27–47). W. B. Saunders. 10.1016/B978-1-4377-0326-9.00003-8

Borbély, A. A., Daan, S., Wirz-Justice, A., & Deboer, T. (2016). The two-process model of sleep regulation: A reappraisal. Journal of Sleep Research, 25(2), 131–143. 10.1111/jsr.12371

Bornkessel-Schlesewsky, I., Sharrad, I., Howlett, C. A., Alday, P. M., Corcoran, A. W., Bellan, V., Wilkinson, E., Kliegl, R., Lewis, R. L., Small, S. L., & Schlesewsky, M. (2022). Rapid adaptation of predictive models during language comprehension: Aperiodic EEG slope, individual alpha frequency and idea density modulate individual differences in real-time model updating. Frontiers in Psychology, 13. 10.3389/fpsyg.2022.817516

Born, J., & Feld, G. B. (2012). Sleep to upscale, sleep to downscale: balancing homeostasis and plasticity. Neuron, 75(6), 933–935. 10.1016/j.neuron.2012.09.007

Brehm, L., & Alday, P. M. (2022). Contrast coding choices in a decade of mixed models. Journal of Memory and Language, 125, 104334. 10.1016/j.jml.2022.104334

Bridger, E. K., Bader, R., Kriukova, O., Unger, K., & Mecklinger, A. (2012). The FN400 is functionally distinct from the N400. NeuroImage, 63(3), 1334–1342. 10.1016/j.neuroimage.2012.07.047

Buysse, D. J., Reynolds, C. F., Monk, T. H., Berman, S. R., & Kupfer, D. J. (1989). The Pittsburgh Sleep Quality Index: A new instrument for psychiatric practice and research. Psychiatry Research, 28(2), 193–213. 10.1016/0165-1781(89)90047-4

Caldwell, H. B., Chatburn, A., Lushington, K., Hanslmayr, S., & Michelmann, S. (2026). Phase similarity between similar objects indicates representational merging across retrieval training but not sleep. Imaging Neuroscience, 4. 10.1162/imag.a.1321

Chatburn, A., Lushington, K., & Cross, Z. R. (2024). Considerations towards a neurobiologically-informed EEG measurement of sleepiness. Brain Research, 1841, 149088. 10.1016/j.brainres.2024.149088

Cross, Z. R., Corcoran, A. W., Schlesewsky, M., Kohler, M. J., & Bornkessel-Schlesewsky, I. (2022). Oscillatory and aperiodic neural activity jointly predict language learning. Journal of Cognitive Neuroscience, 34(9), 1630–1649. 10.1162/jocn_a_01878

Cross, Z. R., Helfrich, R. F., Corcoran, A. W., Dede, A. J. O., Kohler, M. J., Coussens, S. W., Zou-Williams, L., Schlesewsky, M., Gaskell, G. M., Knight, R. T., & Bornkessel-Schlesewsky, I. (2024). Slow Oscillation–Spindle Coupling Predicts Sequence-Based Language Learning. The Journal of Neuroscience, 45(3), e2193232024. 10.1523/jneurosci.2193-23.2024

Dave, S., Brothers, T. A., & Swaab, T. Y. (2018). 1/*f* neural noise and electrophysiological indices of contextual prediction in aging. Brain Research, 1691, 34–43. 10.1016/j.brainres.2018.04.007

Donoghue, T., Haller, M., Peterson, E. J., Varma, P., Sebastian, P., Gao, R., Noto, T., Lara, A. H., Wallis, J. D., Knight, R. T., Shestyuk, A., & Voytek, B. (2020). Parameterizing neural power spectra into periodic and aperiodic components. Nature Neuroscience, 23(12), 1655–1665. 10.1038/s41593-020-00744-x

Fox, J., & Weisberg, S. (2019). An R companion to applied regression (3rd ed.). SAGE.

Gao, R., Peterson, E. J., & Voytek, B. (2017). Inferring synaptic excitation/inhibition balance from field potentials. NeuroImage, 158, 70–78. 10.1016/j.neuroimage.2017.06.078

Gisiger, T. (2001). Scale invariance in biology: coincidence or footprint of a universal mechanism? Biological Reviews, 76(2), 161–209. 10.1017/s1464793101005607

Gramfort, A., Luessi, M., Larson, E., Engemann, D. A., Strohmeier, D., Brodbeck, C., Goj, R., Jas, M., Brooks, T., Parkkonen, L., & Hämäläinen, M. S. (2013). MEG and EEG data analysis with MNE-Python. Frontiers in Neuroscience, 7, Article 267. 10.3389/fnins.2013.00267

He, B. J., Zempel, J. M., Snyder, A. Z., & Raichle, M. E. (2010). The Temporal Structures and Functional Significance of Scale-free Brain Activity. Neuron, 66(3), 353–369. 10.1016/j.neuron.2010.04.020

Helfrich, R. F., Lendner, J. D., & Knight, R. T. (2021). Aperiodic sleep networks promote memory consolidation. Trends in Cognitive Sciences, 25(8), 648–659. 10.1016/j.tics.2021.04.009

Hodapp, A., & Rabovsky, M. (2021). The N400 ERP component reflects an error-based implicit learning signal during language comprehension. European Journal of Neuroscience, 54(9), 7125–7140. 10.1111/ejn.15462

Horne, J. A., & Östberg, O. (1976). A self-assessment questionnaire to determine morningness-eveningness in human circadian rhythms. International Journal of Chronobiology, 4, 97–110.

Hubbard, R. J., Rommers, J., Jacobs, C. L., & Federmeier, K. D. (2019). Downstream behavioral and electrophysiological consequences of word prediction on recognition memory. Frontiers in Human Neuroscience, 13, Article 291. 10.3389/fnhum.2019.00291

Huber, R., Mäki, H., Rosanova, M., Casarotto, S., Canali, P., Casali, A. G., Tononi, G., & Massimini, M. (2013). Human cortical excitability increases with time awake. Cerebral Cortex, 23(2), 1–7. 10.1093/cercor/bhs014

Immink, M. A., Cross, Z. R., Chatburn, A., Baumeister, J., Schlesewsky, M., & Bornkessel-Schlesewsky, I. (2021). Resting-state aperiodic neural dynamics predict individual differences in visuomotor performance and learning. Human Movement Science, 78, 102829. 10.1016/j.humov.2021.102829

Jano, S., Chatburn, A., Cross, Z. R., Schlesewsky, M., & Bornkessel-Schlesewsky, I. (2024). How predictability and individual alpha frequency shape memory: Insights from an event-related potential investigation. Neurobiology of Learning and Memory, 216, Article 108006. 10.1016/j.nlm.2024.108006

Jas, M., Engemann, D. A., Bekhti, Y., Raimondo, F., & Gramfort, A. (2017). Autoreject: Automated artifact rejection for MEG and EEG data. NeuroImage, 159, 417–429. 10.1016/j.neuroimage.2017.06.030

Kuperberg, G. R., Brothers, T., & Wlotko, E. W. (2020). A Tale of Two Positivities and the N400: Distinct Neural Signatures Are Evoked by Confirmed and Violated Predictions at Different Levels of Representation. Journal of Cognitive Neuroscience, 32(1), 12–35. 10.1162/jocn_a_01465

Kuznetsova, A., Brockhoff, P. B., & Christensen, R. H. B. (2017). *lmerTest* package: Tests in linear mixed effects models. Journal of Statistical Software, 82(13), 1–26. 10.18637/jss.v082.i13

Lendner, J. D., Mander, B. A., Schuh-Hofer, S., Schmidt, H., Knight, R. T., Walker, M. P., Lin, J., & Helfrich, R. F. (2022). Human REM sleep controls neural excitability in support of memory formation. bioRxiv. 10.1101/2022.05.13.491801

Litwin-Kumar, A., & Doiron, B. (2014). Formation and maintenance of neuronal assemblies through synaptic plasticity. Nature Communications, 5(1), Article 1. 10.1038/ncomms6319

Lo, J. C., Dijk, D.-J., & Groeger, J. A. (2014). Comparing the Effects of Nocturnal Sleep and Daytime Napping on Declarative Memory Consolidation. PLoS ONE, 9(9), e108100. 10.1371/journal.pone.0108100

Makowski, D. (2018). The *psycho* package: An efficient and publishing-oriented workflow for psychological science. Journal of Open Source Software, 3(22), 470. 10.21105/joss.00470

MacGregor-Fors, I., & Payton, M. E. (2013). Contrasting Diversity Values: Statistical Inferences Based on Overlapping Confidence Intervals. PLoS ONE, 8(2), e56794. 10.1371/journal.pone.0056794

Mathôt, S., Schreij, D., & Theeuwes, J. (2012). OpenSesame: An open-source, graphical experiment builder for the social sciences. Behavior Research Methods, 44(2), 314–324. 10.3758/s13428-011-0168-7

Nere, A., Hashmi, A., Cirelli, C., & Tononi, G. (2013). Sleep-dependent synaptic down-selection (I): Modeling the benefits of sleep on memory consolidation and integration. Frontiers in Neurology, 4, Article 143. 10.3389/fneur.2013.00143

Nicholls, M. E. R., Thomas, N. A., Loetscher, T., & Grimshaw, G. M. (2013). The Flinders Handedness survey (FLANDERS): A brief measure of skilled hand preference. Cortex: A Journal Devoted to the Study of the Nervous System and Behavior, 49, 2914–2926. 10.1016/j.cortex.2013.02.002

Niethard, N., Burgalossi, A., & Born, J. (2017). Plasticity during sleep is linked to specific regulation of cortical circuit activity. Frontiers in Neural Circuits, 11, Article 65. 10.3389/fncir.2017.00065

Nishida, M., & Walker, M. P. (2007). Daytime Naps, Motor Memory Consolidation and Regionally Specific Sleep Spindles. PLoS ONE, 2(4), e341–e341. 10.1371/journal.pone.0000341

Ognjanovski, N., Schaeffer, S., Wu, J., Mofakham, S., Maruyama, D., Zochowski, M., & Aton, S. J. (2017). Parvalbumin-expressing interneurons coordinate hippocampal network dynamics required for memory consolidation. Nature Communications, 8(1), Article 15039. 10.1038/ncomms15039

Olcese, U., Esser, S. K., & Tononi, G. (2010). Sleep and synaptic renormalization: A computational study. Journal of Neurophysiology, 104(6), 3476–3493.

Ouyang, G., Hildebrandt, A., Schmitz, F., & Herrmann, C. S. (2020). Decomposing alpha and 1/f brain activities reveals their differential associations with cognitive processing speed. NeuroImage, 205, 116304. 10.1016/j.neuroimage.2019.116304

Payne, J. D., Kensinger, E. A., Wamsley, E. J., Spreng, R. N., Alger, S. E., Gibler, K., Schacter, D. L., & Stickgold, R. (2015). Napping and the selective consolidation of negative aspects of scenes. Emotion, 15(2), 176–186. 10.1037/a0038683

Rabovsky, M., Hansen, S. S., & McClelland, J. L. (2018). Modelling the N400 brain potential as change in a probabilistic representation of meaning. Nature Human Behaviour, 2(9), 693–705. 10.1038/s41562-018-0406-4

Rasch, B., & Born, J. (2013). About sleep’s role in memory. Physiological Reviews, 93(2), 681–766. 10.1152/physrev.00032.2012

Sassenhagen, J., & Alday, P. M. (2016). A common misapplication of statistical inference: Nuisance control with null-hypothesis significance tests. Brain and Language, 162, 42–45. 10.1016/j.bandl.2016.08.001

Stark, S. M., Kirwan, C. B., & Stark, C. E. L. (2019). Mnemonic Similarity Task: A tool for assessing hippocampal integrity. Trends in Cognitive Sciences, 23(11), 938–951. 10.1016/j.tics.2019.08.003

Stickgold, R. (2009). How do I remember? Let me count the ways. Sleep Medicine Reviews, 13(5), 305–308. 10.1016/j.smrv.2009.05.004

Stróżak, P., Abedzadeh, D., & Curran, T. (2016). Separating the FN400 and N400 potentials across recognition memory experiments. Brain Research, 1635, 41–60. 10.1016/j.brainres.2016.01.015

Tononi, G., & Cirelli, C. (2003). Sleep and synaptic homeostasis: A hypothesis. Brain Research Bulletin, 62(2), 143–150. 10.1016/j.brainresbull.2003.09.004

Tononi, G., & Cirelli, C. (2006). Sleep function and synaptic homeostasis. Sleep Medicine Reviews, 10(1), 49–62. 10.1016/j.smrv.2005.05.002

Tononi, G., & Cirelli, C. (2012). Time to Be SHY? Some comments on sleep and synaptic homeostasis. Neural Plasticity, 2012, e415250. 10.1155/2012/415250

Tononi, G., & Cirelli, C. (2014). Sleep and the price of plasticity: From synaptic and cellular homeostasis to memory consolidation and integration. Neuron, 81(1), 12–34. 10.1016/j.neuron.2013.12.025

Vallat, R., & Walker, M. P. (2021). An open-source, high-performance tool for automated sleep staging. eLife, 10, e70092. 10.7554/eLife.70092

Vyazovskiy, V. V. (2013). Cortical neuronal mechanisms of sleep homeostasis. Zhurnal Vysshei Nervnoi Deiatelnosti Imeni I P Pavlova, 63(1), 13–23. 10.7868/s0044467713010176

Vyazovskiy, V. V., Cirelli, C., Pfister-Genskow, M., Faraguna, U., & Tononi, G. (2008). Molecular and electrophysiological evidence for net synaptic potentiation in wake and depression in sleep. Nature Neuroscience, 11(2), Article 2. 10.1038/nn2035

Vyazovskiy, V. V., Olcese, U., Lazimy, Y. M., Faraguna, U., Esser, S. K., Williams, J. C., Cirelli, C., & Tononi, G. (2009). Cortical firing and sleep homeostasis. Neuron, 63(6), Article 6. 10.1016/j.neuron.2009.08.024

Walker, M. P., & Stickgold, R. (2004). Sleep-dependent learning and memory consolidation. Neuron, 44(1), 121–133. 10.1016/j.neuron.2004.08.031

Walker, M. P., & Stickgold, R. (2006). Sleep, memory, and plasticity. Annual Review of Psychology, 57(1), 139–166. 10.1146/annurev.psych.56.091103.070307

Warriner, A. B., Kuperman, V., & Brysbaert, M. (2013). Norms of valence, arousal, and dominance for 13,915 English lemmas. Behavior Research Methods, 45(4), 1191–1207. 10.3758/s13428-012-0314-x

Wen, H., & Liu, Z. (2016). Separating fractal and oscillatory components in the power spectrum of neurophysiological signal. Brain Topography, 29(1), 13–26. 10.1007/s10548-015-0448-0

Weymar, M., Löw, A., Modess, C., Engel, G., Gründling, M., Petersmann, A., Siegmund, W., & Hamm, A. O. (2010). Propranolol selectively blocks the enhanced parietal old/new effect during long-term recollection of unpleasant pictures: A high density ERP study. NeuroImage, 49, 2800–2806. 10.1016/j.neuroimage.2009.10.025

Wickham, H. (2016). ggplot2: Elegant graphics for data analysis. Springer. https://ggplot2.tidyverse.org

